# Moving pictures: Reassessing docking experiments with a dynamic view of protein interfaces

**DOI:** 10.1101/2020.12.08.415885

**Authors:** Chantal Prévost, Sophie Sacquin-Mora

**Affiliations:** CNRS, Laboratoire de Biochimie Théorique, UPR9080, Université de Paris, 13 rue Pierre et Marie Curie, 75005 Paris, France; Institut de Biologie Physico-Chimique, Fondation Edmond de Rothschild PSL Research University, 75006 Paris, France

**Keywords:** protein interactions, protein interfaces, docking, molecular dynamics simulations

## Abstract

The modeling of protein assemblies at the atomic level remains a central issue in structural biology, as protein interactions play a key role in numerous cellular processes. This problem is traditionally addressed using docking tools, where the quality of the models is based on their similarity to a single reference experimental structure. However, using a static reference does not take into account the dynamic quality of the protein interface. Here, we used all-atom classical Molecular Dynamics simulations to investigate the stability of the reference interface for three complexes that previously served as targets in the CAPRI competition. For each one of these targets, we also ran MD simulations for ten models that are distributed over the *High*, *Medium* and *Acceptable* accuracy categories. To assess the quality of these models from a dynamic perspective, we set up new criteria which take into account the stability of the reference experimental protein interface. We show that, when the protein interfaces are allowed to evolve along time, the original ranking based on the static CAPRI criteria no longer holds as over 50% of the docking models undergo a category change (which can be either toward a better or a lower accuracy group) when reassessing their quality using dynamic information.

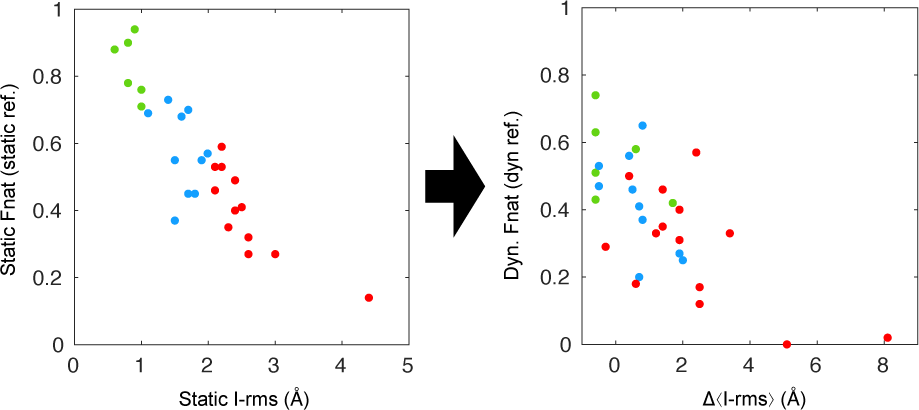

## 1. Introduction

A protein biological activity in the cellular environment heavily relies on its ability to insert itself in a complex interaction network involving several macromolecular partners that can be other proteins or nucleic acids.^1, 2^ As a consequence, deciphering the protein social network remains a key issue in our understanding of the cell function, with important consequences for pharmacological developments.^3–7^ This appeared very clearly during the recent COVID-19 crisis, as hundreds of preprints investigating SARS-CoV-2 protein interactions (and sometimes their inhibition by various drugs) were deposited on repositories such as BioRxiv, thus highlighting the importance of our understanding of protein interactions in order to develop potential cures or vaccines against this virus. From the experimental point of view, proteomic studies provide us with a wealth of information regarding proteins networks both at the cellular and at the whole organism level.^8, 9^ Structural data regarding macromolecular assemblies is available in the Protein Data Bank (PDB),^10^ and the number of protein complexes structures released each year has been steadily increasing over the last decade, in particular thanks to recent advances in cryo-EM techniques, which enabled to reach atomic resolution for very large complexes.^11, 12^ On the other hand, in silico methods represent a complementary approach for the identification of protein partners^13–15^ and determining the 3-dimensional structure of protein assemblies.^16–18^ *Integrative* modeling techniques, which include evolutionary information or experimental data when building a protein complex model are of particular interest, especially when working on more challenging systems (such as flexible assemblies or membrane associated complexes^19^), as they will combine the best of both worlds.^20–22^

A central problem when modeling protein interactions is the *docking* question, where one attempts to predict the atomic structure of a protein complex based on the structure of its individual components. The CAPRI (for Critical Assessment of PRedicted Interactions, https://www.capri-docking.org/) initiative, which was created in 2001 has played a central role in stimulating development and progress in docking and scoring methods.^23–25^ Over the years, dozens of research groups have had the opportunity to test the performance of their computational procedures against the blind prediction of more than 150 *targets*, i.e. macromolecular assemblies for which the experimental structure was provided to CAPRI prior to publication. Since the beginnings of CAPRI, the assessment of the quality of the predictions relies on criteria which evaluate how close the submitted models are to the reference crystallographic structure, i.e. the fraction of native contacts, the ligand and the interface root mean square deviations.^26, 27^ However, these criteria, which are based on a single protein structure, convey a static vision of protein-protein interactions that is now being increasingly questioned. More and more data suggest that protein interfaces are dynamic objects, sometimes including disordered, flexible segments and conformational heterogeneity.^28–33^ This aspect should be taken into account more often,^34–36^ in particular as it plays a part in the specificity of protein interactions.^37–39^ With these questions in mind, we used all-atom classical molecular dynamics simulation to investigate how model protein interfaces will behave along time compared to the original crystallographic interface. This involved redefining new criteria based on the dynamic behavior of the reference structure, and seeing wether the original CAPRI ranking of the docking models into the *High*, *Medium* and *Acceptable* categories still holds, once these model interfaces have also been allowed to evolve along time.

## 2. Material and Methods

### Docking models selection

The CAPRI Score-set (http://cb.iri.univ-lille1.fr/Users/lensink/Score_set/) set up by Lensink and Wodak^40^ lists models produced by participants to the CAPRI competition for 15 protein complexes (targets), with between 300 and 2000 docking poses available for each target. In the present study, we selected the three targets (41, 29 and 37, shown in Figure 1) for which models are available for the three accuracy categories High, Medium and Acceptable (see the Supplementary information for a reminder of the CAPRI assessment criteria defining theses categories). Target 41 (pdb code 2wpt^41^) is a complex between the colicin E9 endonuclease, an antibiotic protein, and its bacterial inhibitor, the immunity protein Im2 from *E. coli*. Target 29 (pdb code 2vdu^42^) is a tRNA m7G methylation complex of yeast, which is involved in the tRNA degradation pathway. It comprises a catalytic unit Trm8 (chain D in the pdb file) bound to the non catalytic unit Trm82 (chain F in the pdb file), which modulates the Trm8 activity. Finally, Target 37 (pdb code 2w83^43^) is a trimeric assembly formed by the human GTP-binding ADP-ribosylation factor ARF6 bound with two leucine zipper domains from the JIP4 protein. This interaction regulates the binding of JIPs with motor proteins such as kinesin or dynactin. These three complexes present buried surfaces areas (BSA) comprised between 1500 and 2000 Å^2^ (see Table 1), which is relatable to the average BSA of 1600 Å that was observed for single patch interfaces by Chakrabarty and Janin in their review of protein protein recognition sites^44^. For each target, ten models were selected that are distributed over the three categories and are listed in Table 1. When possible, models with the lowest number of steric clashes were selected.

**Figure 1:**
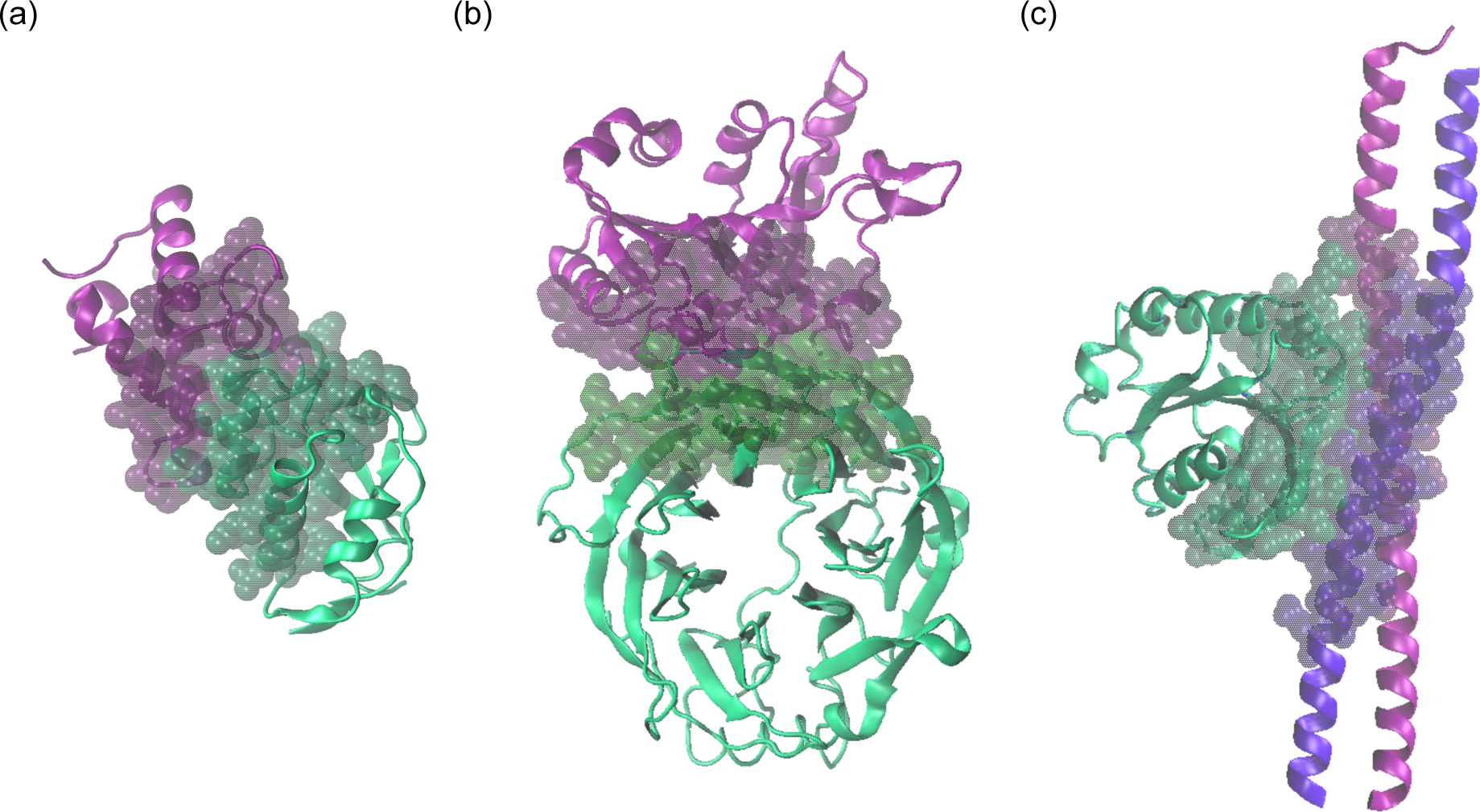
Cartoon representations of the target structures under study with the receptor chain in green and the ligand chain in purple. Residues belonging to the static interface are shown as transparent van der Waals spheres. (a) T41, (b) T29, (c) T37. The images in this figure and Figure 4 were prepared using Visual Molecular Dynamics.^53^

**Table 1.** Models from the CAPRI Score-set used for this study.

| Target | Buried Surface Area (Å<sup>2</sup>) | High models Ids | Medium models Ids | Acceptable models Ids |
| --- | --- | --- | --- | --- |
| **T41 (PDB 2WPT, chains B, A)** | 1566 | 604, 1009 | 31, 241, 1149 | 50, 61, 302, 380, 490 |
| **T29 (PDB 2VDU, chains D,F)** | 1990 | 190, 1851 | 56, 173, 875, 1818 | 608, 1631, 1909, 2109 |
| **T37 (PDB 2W83, chains A, CD)** | 1719 | 304, 852 | 68, 351, 833 | 146, 660, 960, 1022, 1275 |

### Molecular Dynamics Simulations

We performed all-atom classical Molecular Dynamics (MD) simulations for the crystallographic structures of targets 41, 37, and 29 and also for all the selected models from Table 1. We used Gromacs, version 5.0.4, with the OPLS-AA force field^45^ and periodic boundary conditions. The first step of the simulation procedure is an *in vacuo* minimization of the structure with the steepest descent algorithm during 5000 steps without any constraints. We added a triclinic water box of 2 nm around the protein, filled with TIP3P molecule type^46^ and the system was neutralized with the addition of ions randomly placed in the box while maintaining the NaCl concentration at 150 mM. For Target 41/37/29 crystallographic structure, the whole system contains around 3200/5500/9300 atoms, 22500/60000/40000 water molecules and 120/360/250 ions (with approximatively 63/185/132 Na^+^ and 60/175/120 Cl^-^). A second minimization step was performed with the same set of parameters as before during 5000 steps to prevent possible water clashes with the protein. Molecular Dynamics Simulations were carried out using an integration time step of 2 fs. Initial heating and pre-equilibration steps were carried out by assigning random velocities from the Maxwell-Boltzmann distribution, performing 100 ps of dynamics under NVT conditions, and then 100 ps under NPT conditions. During this process and for all subsequent simulations the temperature was fixed at 300 K using the velocity rescale method.^47^ All covalent bonds involving hydrogen atoms were constrained with the LINCS algorithm^48^ and electrostatic interactions were computed using the Particle Mesh Ewald method^49^. For the pressure coupling during the NPT equilibration, we used the Parinello-Rahman method^50^ at the value of 1 atm. Production phases were finally done using the same set of parameters and algorithms during 100 ns. Trajectories were saved every 10 ps, and we used 1000 frames (one frame every 100 ps) to calculate average values (listed below) over the trajectory.

### Interfaces analysis

The CAPRI assessment criteria^40^ (listed in table SI-1) are based on three parameters :

- *f_nat_* is the fraction of receptor-ligand residue contacts in the target structure that are reproduced in the model (residues are considered to be in contact if any of their heavy atoms are within 5 Å).

- L-rms is the root mean square deviation (rmsd) for the ligand protein backbone atoms with respect to the reference structure.

- I-rms is the rmsd for the interface backbone atoms. Interface residues are defined using a 10 Å distance threshold between the interaction partners (with atom-atom contacts below 3 Å defined as clashes).

These three parameters use the protein complex experimental structure as a reference, and will be referred to as *static* parameters for the rest of the study. Figures 2a-b display the distribution of these parameters for the 30 models under study and highlights the separation of the High, Medium and Acceptable groups.

**Figure 2:**
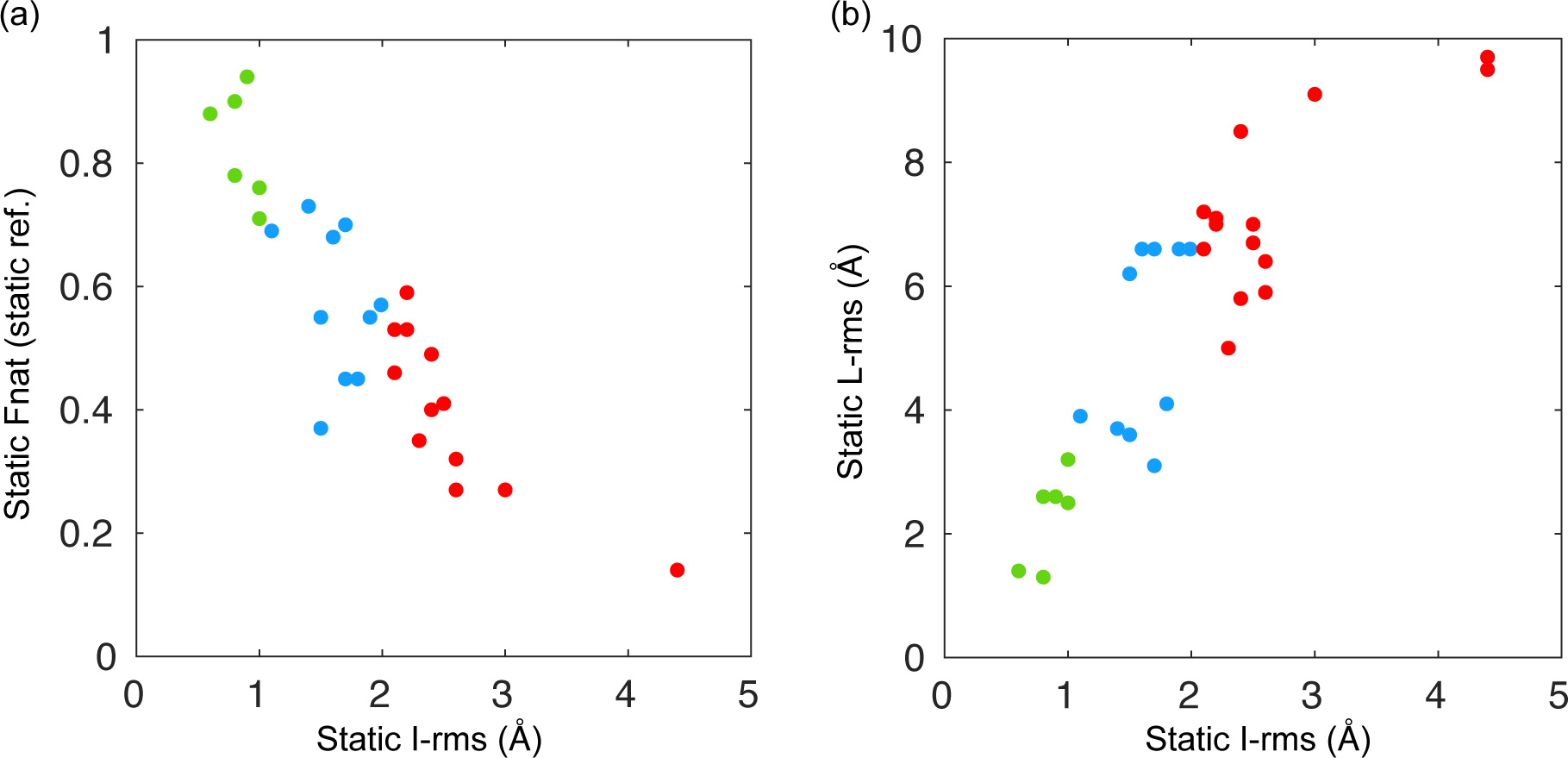
(a) Static *f_nat_* as a function of I-rms (b) L-rms as a function of I-rms. Green: High quality models. Blue: Medium quality models. Red: Acceptable quality models. This color code will be used all throughout the manuscript.

In order to take into account changes in the protein interface that might occur during the MD simulations, we defined the following *dynamic* parameters:

- *Dynamic contacts* concern residues that are in contact during at least 50% of the trajectory (that is, 50% of the 1000 frames kept for calculating average values). As a consequence *f_nat_* can now be computed by using static contacts both in the model and the experimental structure (*f_nat-statref_*), or by using dynamic contacts in the model (calculated over the whole trajectory) and comparing them to static (*dyn-f_nat-statref_*), or dynamic (*dyn-f_nat-dynref_*), contacts in the experimental structure. For the remainder of the study, the term *dyn-f_nat_* will refer to the fraction of native contacts when comparing dynamic contacts in the model and in the reference (*dyn-f_nat-dynref_*).

- Instead of the L-rms, we can consider its average value ⟨L-rms⟩ over the whole trajectory, for simulations starting from either the reference structure, or for a model, where L-rms’s used for calculating the average are takesn with respect to the (static) reference structure.

- The *dynamic interface* is defined as residues from the protein partners that are less than 10 Å away during at least 50% of the trajectory. As a consequence the average value for the I-rms can be calculated using either the static interface residues or the dynamic interface residues

All the values for the static and dynamic parameters listed above and obtained after performing molecular dynamics simulations for the crystallographic structures and the 30 models are listed in Table SI-2 in the supplementary information.

## 3. Results and discussion

### Stability of the reference interfaces

The stability of the crystallographic reference interfaces along time was investigated by monitoring the L-rms, I-rms and *f_nat_* values during the trajectories (see Figure 3), and Figure SI-1 presents snapshots of the complexes showing the position of the ligand chains by the end of the 100 ns trajectory. While the L-rms and I-rms values quickly leave the high-quality area (highlighted in green) at the beginning of the simulations, they remain in the medium-acceptable area (highlighted in blue-red) for most of the trajectory (see Figure 3). In the case of Target 37 (pdb 2w83), the high values observed for the L-rms after 20 ns are due to the rotation of the long ⍺-helices forming the ligand chains and their deformation (see the kink in helix C, in blue, in Figure SI-1c). However, these deformations are limited to the extremities of the helices, and the protein interface remains stable, as shown from the I-rms value on Figure 3h. Meanwhile, the *f_nat_* values remain in the [0.6-0.8] range during the complete simulation for all three targets (see Figures 3c,f,i). A similar behavior was already observed in earlier modeling studies on the monomer-monomer interface in the RecA filament,^51^ which supports the idea that no destabilization of the reference crystallographic interfaces occurs during the MD simulations. In the case of Target 29 (pdb 2vdu), one can even observe an increase in the contact number after 40 ns (Figure SI-3a), which is due to the N-terminal and C-terminal tails of the ligand protein forming new contacts with the surface of the receptor protein (see Figure SI-1b). Finally, one can map the contact residues over the protein surfaces for three targets. Figure 4 shows the contact residues in the reference interfaces (in green or purple, for the receptor or ligand protein respectively), or colored as the fraction of the trajectory during which they remain in contact with the interaction partner (white, no contact during the trajectory, green or purple, the contact lasts for the whole trajectory). Once again, one can see how the protein interfaces remain stable, with a strong conservation of the central patch, and small variations on the external borders (thus reminding us of the rim/core classification of protein interface residues set up by Levy^52^).

**Figure 3:**
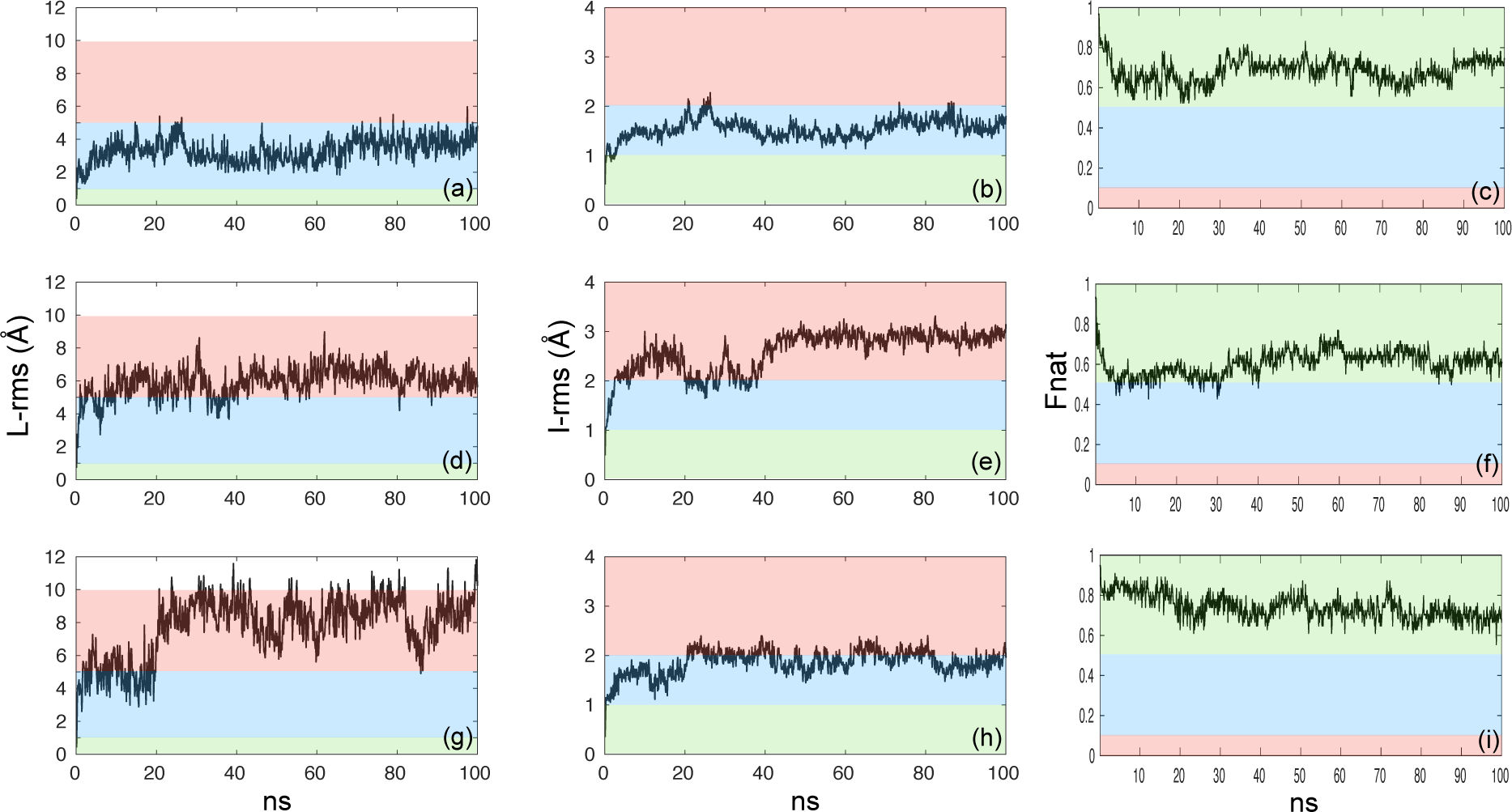
Time evolution of the L-rms, I-rms and f_nat_ values for the three crystallographic reference interfaces : T41 (a), (b), (c); T29 (d), (e), (f); T37 (g), (h), (i). The areas corresponding to the high, medium and acceptable categories are highlighted with green, blue and red backgrounds respectively.

**Figure 4:**
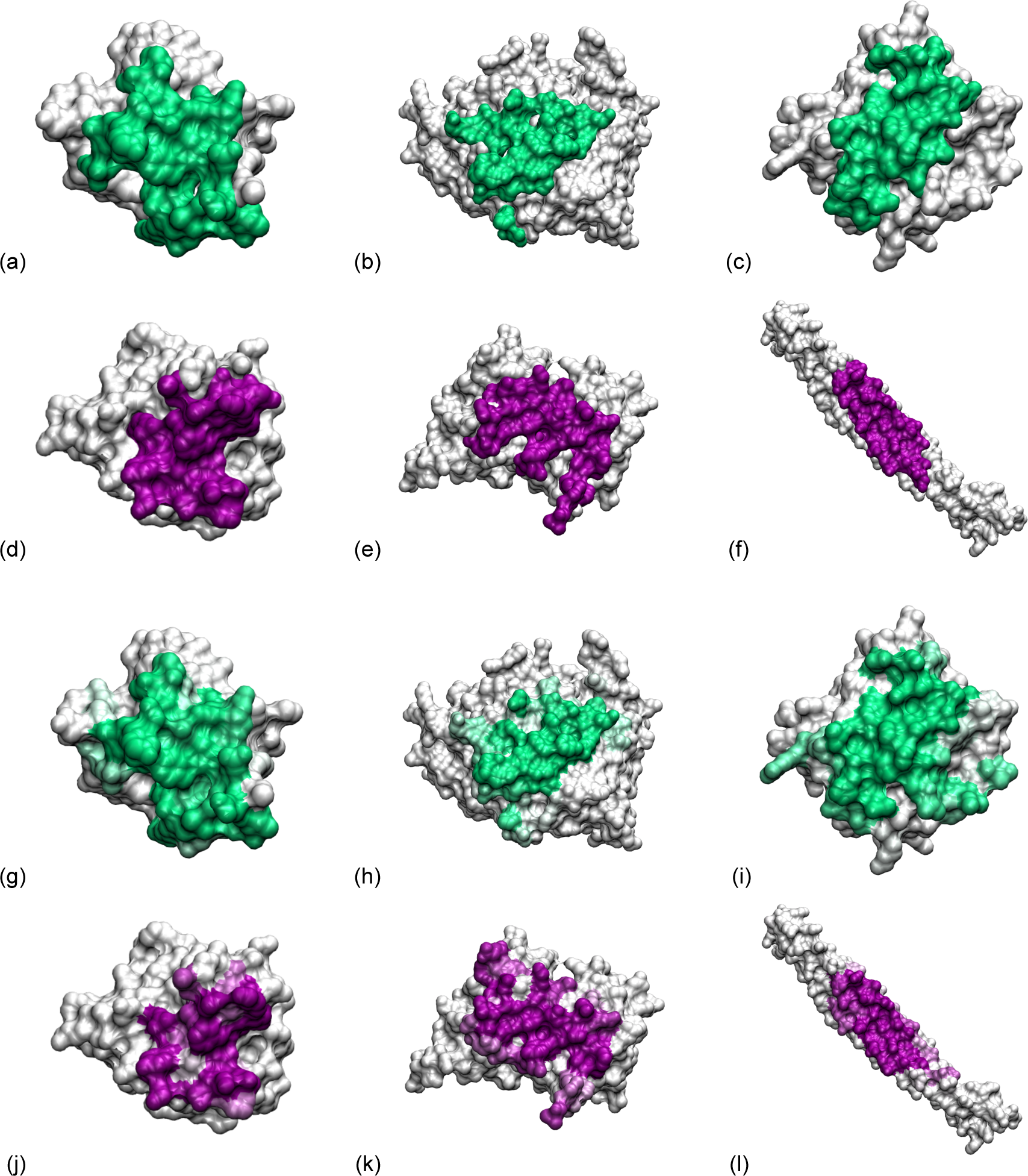
Mapping the contact residues on the protein surface. Contacts residues in the reference interface shown in green and purple for the receptor and ligand protein respectively : (a) Target 41 chain A, (b) Target 29 chain D, (c) Target 37 chain A, (d) Target 41 chain B, (e) Target 29 chain F, (f) Target 37 chains CD. Residues colored as the fraction of the trajectory during which they remain in contact with the interaction partner (white, no contact during the trajectory, green or purple, the contact lasts for the whole trajectory) : (g) Target 41 chain A, (h) Target 29 chain D, (i) Target 37 chain A, (j) Target 41 chain B, (k) Target 29 chain F, (l) Target 37 chains CD.

### Residue contacts

Figure 5a shows the evolution of the contacts numbers when going from the static to the dynamic definition, which, on average, leads to a slight decrease (roughly 20% over the 33 MD trajectories, see columns 3 and 4 in Table SI-2). In Figure 5b we compare the *f_nat_* value using the static and dynamic definitions. The horizontal and vertical dashed lines highlight the frontiers between the high (*f_nat_* > 0.5), medium (*f_nat_* > 0.3) and acceptable categories (*f_nat_* > 0.1). Interestingly, a large number of models might belong to a different category when using a dynamic definition for the native contacts instead of the static definition. For example, high quality models H1851T29 and H604T41 would now be classified as medium, since their *dyn-f_nat_* value falls below 0.5. On the other hand, the acceptable models A960T37 and A1257T37 see an increase of their *dyn- f_nat_* value, which might lead to enter the medium or high groups (depending on how the L-rms and I-rms values evolved, see below). Note that only roughly one third of the residue contacts that are formed during the MD trajectories are stable enough to be considered as dynamic contacts. Most residue-residue contacts are transient ones that form during 10% or less of the simulation (see Figure SI-5 for the distribution of the contacts times for the three crystallographic structures).

**Figure 5:**
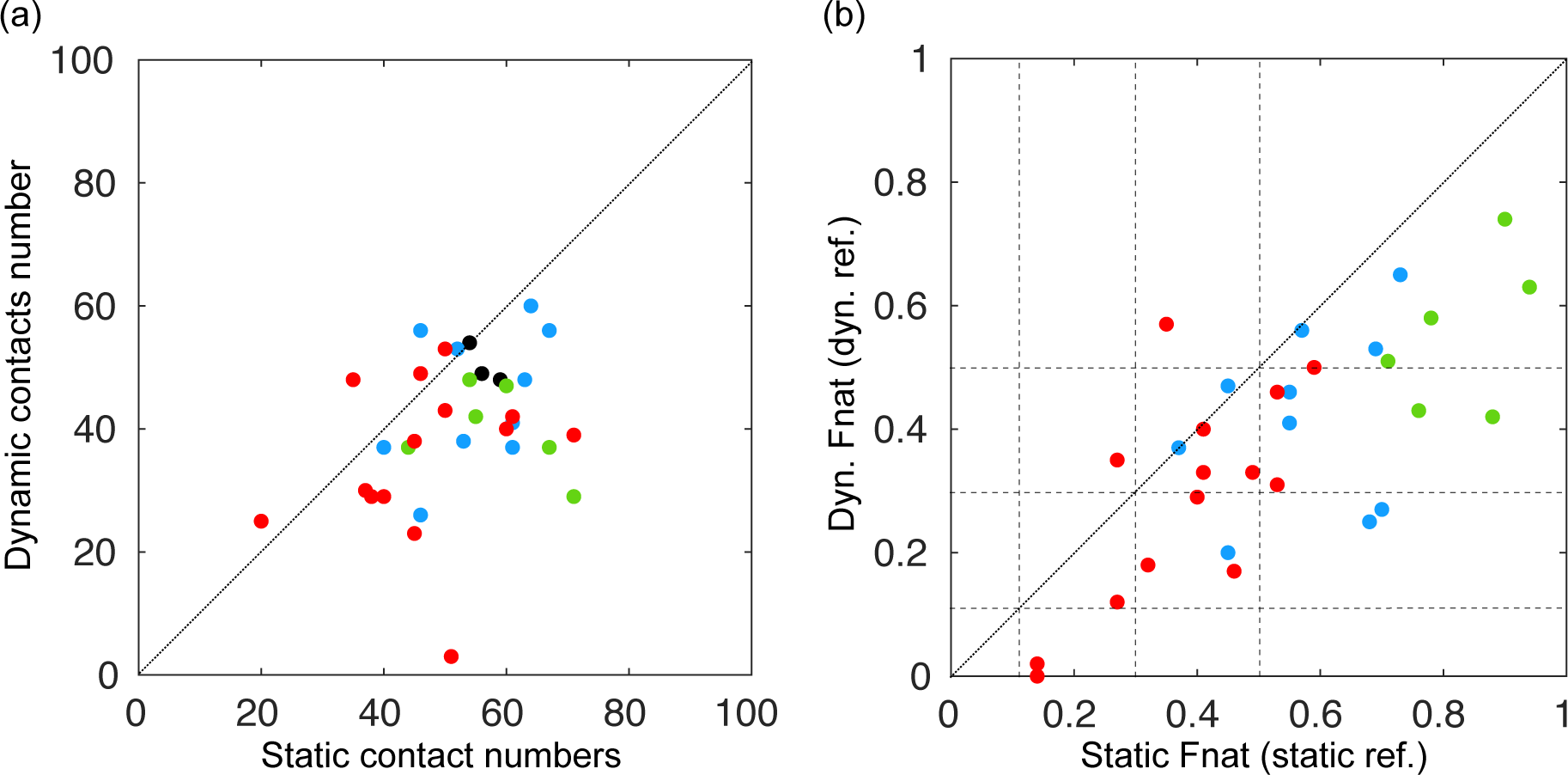
(a) Dynamic contacts number as a function of the static contacts number. The three black dots correspond to the target crystallographic structures, the dotted diagonal is plotted as a guide for the eye. (b) *Dyn-f_nat_* as a function of the static *f_nat_*. The horizontal and vertical dashed lines highlight the frontiers between the high, medium and acceptable categories, and the dotted diagonal is plotted as a guide for the eye.

### L-rms evolution

#### About steric clashes

For each target the few high quality models displayed an important number of steric clashes (between 8 and 18 clashes), while for the medium and acceptables models, we were able to select structures with a lower number of clashes. As a consequence, we first checked that, when present, initial clashes have no visible impact on the stability of the protein complex during the MD simulations. Figure SI-6 shows the L-rms value after the equilibration steps and before production as a function of the number of clashes in the initial structure (the raw data is available in column 2 of Table SI-2). The L-rms is always below 2.0 Å and does not depend on *n_clash_* at all.

As could be expected, the average ⟨L-rms⟩ over the whole MD trajectory is almost always larger than the L-rms in the initial model structure (see Figure 6a). However, for most models the ⟨L-rms⟩ value is of the same order as the average ⟨L-rms⟩cryst observed when using the crystallographic target structures as a starting point (see the black dots on Figure 6a). On Figure 6b, we now compare the L-rms in the initial model structure and the difference between the ⟨L-rms⟩ value obtained for the model and for the corresponding crystallographic target structure (Δ⟨L-rms⟩). Again, the horizontal and vertical dashed lines highlight the frontiers between the high (L-rms < 1.0 Å, no model fulfills this criterion in the initial structure), medium (L-rms < 5.0 Å) and acceptable categories (L-rms < 10.0 Å). A few acceptable models (notably A1022T37 and A608T29, with Δ⟨L-rms⟩ of 23.4 and 12.7 Å respectively), and a high quality one (H604T41, with Δ⟨L-rms⟩ = 6.6 Å) display large Δ⟨L-rms⟩ values, as their structure slowly drifts away from the reference during the simulation. But numerous points also lie below the diagonal, some even showing negative Δ⟨L-rms⟩ values, thus suggesting that the model structure more stably samples the conformational subspace associated to the reference structure than the reference structure itself, even though this model originally belongs to the medium or acceptable category (for example models A1909T37 and A2109T37).

**Figure 6:**
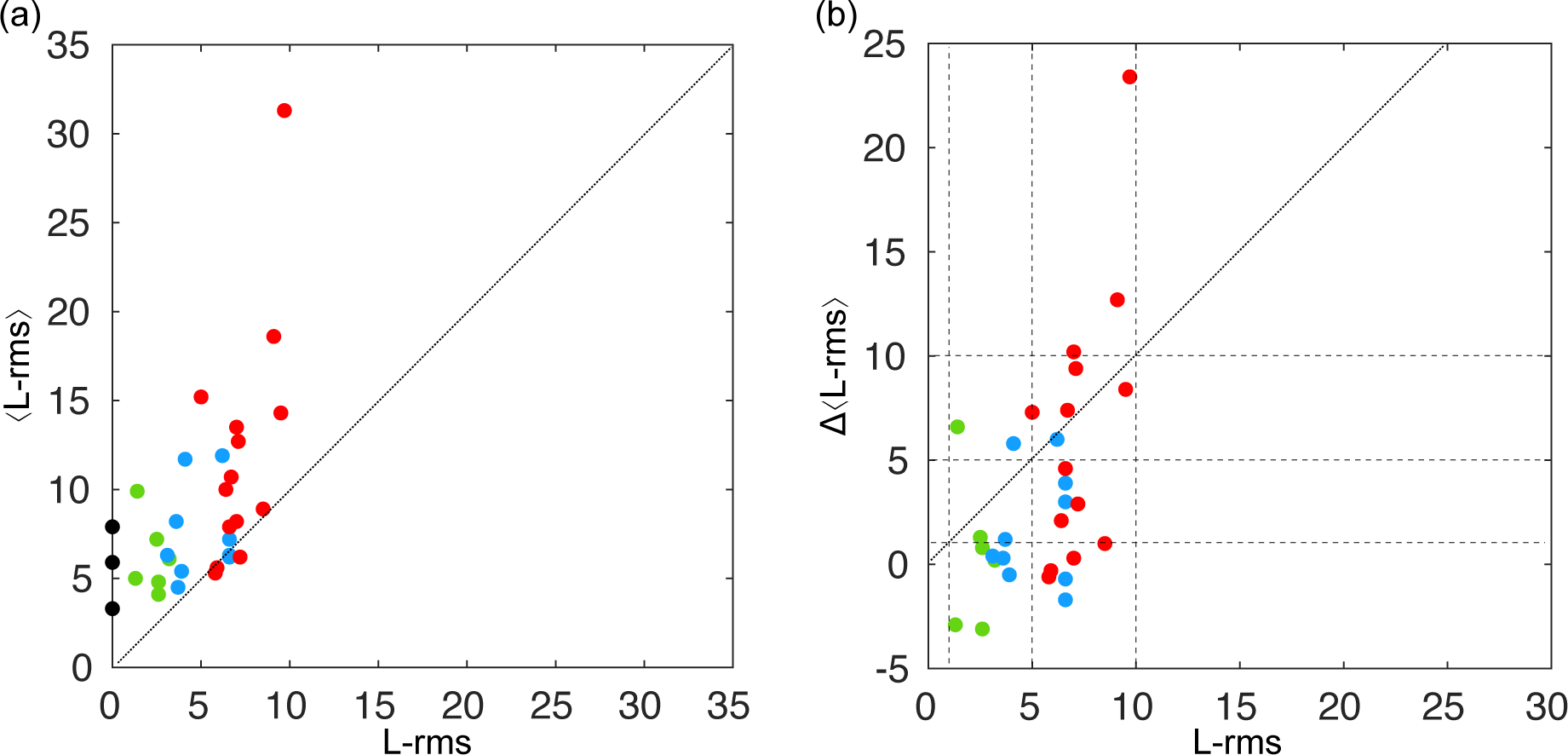
(a) Average L-rms value over the MD trajectory as a function of the L-rms value for the model structure (green, blue and red dots) or the crystallographic structures (black dots). (b) difference between the ⟨L-rms⟩ value obtained for the model and for the corresponding crystallographic target structure (Δ⟨L-rms⟩) as a function of the L-rms value for the model structure. The horizontal and vertical dashed lines highlight the frontiers between the high, medium and acceptable categories, and the dotted diagonal is plotted as a guide for the eye.

### Evolution of the interfaces

For each target we can now define a static reference interface (SRI) based on the crystallographic structure, and a dynamic reference interface (DRI) based on residues that are less than 10 Å apart during at least 50% of the MD simulation using the crystallographic structure as a starting point. For each target, the residues forming the SRI and DRI are listed in Table SI-3 (and SRI residues are shown as van der Waals spheres on Figure 1). Due to the large criterion used for defining the interface residues, these do not vary much in the static and dynamic interfaces, and the average ⟨I-rms⟩ over the MD trajectory almost does not change when calculated with the SRI or the DRI (see Table SI-2, columns 13-16). In a similar fashion to what we did for the L-rms, Figure 7 shows both the ⟨I-rms⟩ and the Δ⟨I-rms⟩ as a function of the initial I-rms in the model on the left and right panels respectively. Again, a large number of points lie below the diagonal, with medium and acceptable models that actually spend more time close to the reference structure than the reference structure itself, and present negative Δ⟨I-rms⟩ values (for example models M68T37 or A2019T29). On the opposite side of the diagonal, the structure of the high quality model H604 for target 41 drifts away from its initial pose during the MD trajectory.

**Figure 7:**
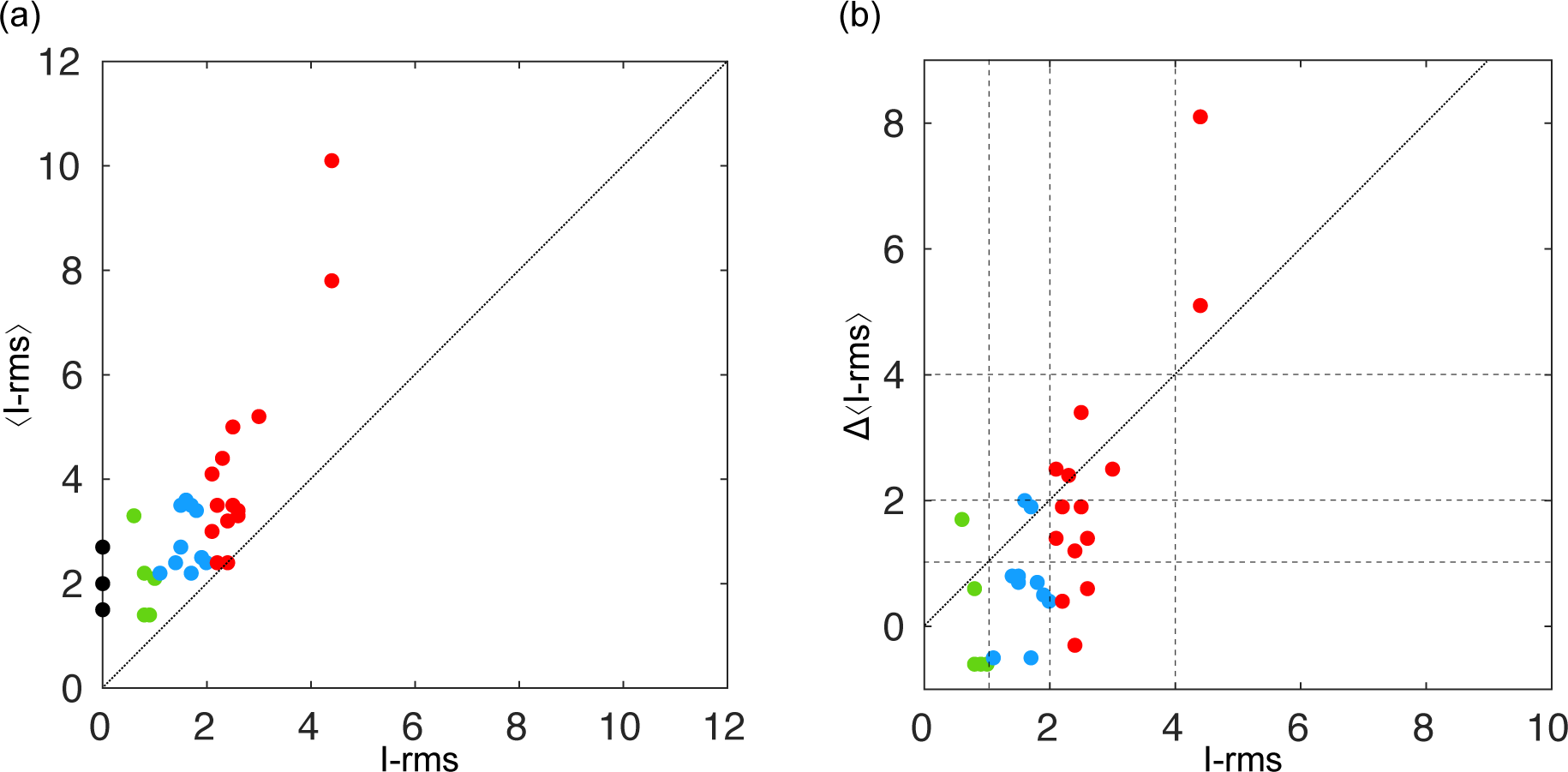
(a) Average I-rms value over the MD trajectory as a function of the I-rms value for the model structure (green, blue and red dots) or the crystallographic structures (black dots). (b) difference between the ⟨I-rms⟩ value obtained for the model and for the corresponding crystallographic target structure (Δ⟨I-rms⟩) as a function of the I-rms value for the model structure. The horizontal and vertical dashed lines highlight the frontiers between the high, medium and acceptable categories, and the dotted diagonal is plotted as a guide for the eye.

### Using dynamic criteria for the ranking of complex models ?

Following these results, it appears that the original, static, CAPRI criteria for ranking protein-protein complex might not hold when switching to a dynamic view of the protein interface. This is especially true for the High category, since not even the original reference experimental structures of the targets display average ⟨L-rms⟩ and ⟨I-rms⟩ values below 1 Å (cf. the black dots on Figures 6a and 7a). One should also note that the *L-rms ≤ 1 Å* criterion used to classify a model in the High category appears to be extremely stringent, and was actually never fulfilled by any of the selected docking models. The six models belonging to the High category in our study only fulfill the I*-rms ≤ 1 Å* criterion but they all display L-rms values above 1 Å (both from a static or a dynamic point of view).

In our dynamic perspective, one could consider that a good quality model is not only structurally close to the original crystallographic structure, but that it also has to be as stable along time as the crystallographic structure. And this stability can be measured via the Δ⟨L-rms⟩ and Δ⟨I-rms⟩ parameters. So, in a similar fashion to what we did in Figure 2, the distribution of the dynamic parameters, *dyn-f_nat_*, Δ⟨L-rms⟩ and Δ⟨I-rms⟩ is shown on Figure 8. The original model categories, that were based on the static parameters values, no longer hold, and we can see that the points for models belonging to different categories are not separated anymore. The areas with colored background in Figures 8ab highlight the high, medium and acceptable accuracy categories when using dynamic parameters (*dyn-f_nat_*, Δ⟨L-rms⟩ and Δ⟨I-rms⟩) instead of static ones (*f_nat_*, L-rms and I-rms) to apply the CAPRI criteria. Numerous models will undergo a category change, and these are summarized in Table 2. Altogether, only 14 out of the 30 models stay in their original category when we consider their dynamic behavior, and the 16 category changes that were observed are equally distributed between 8 upgrades toward a better category and 8 downgrades toward a lower one. Two acceptable models, A1631T29 and A1022T37, eventually end up as incorrect models as their fraction of native contacts drops down to zero during the MD trajectories.

**Figure 8:**
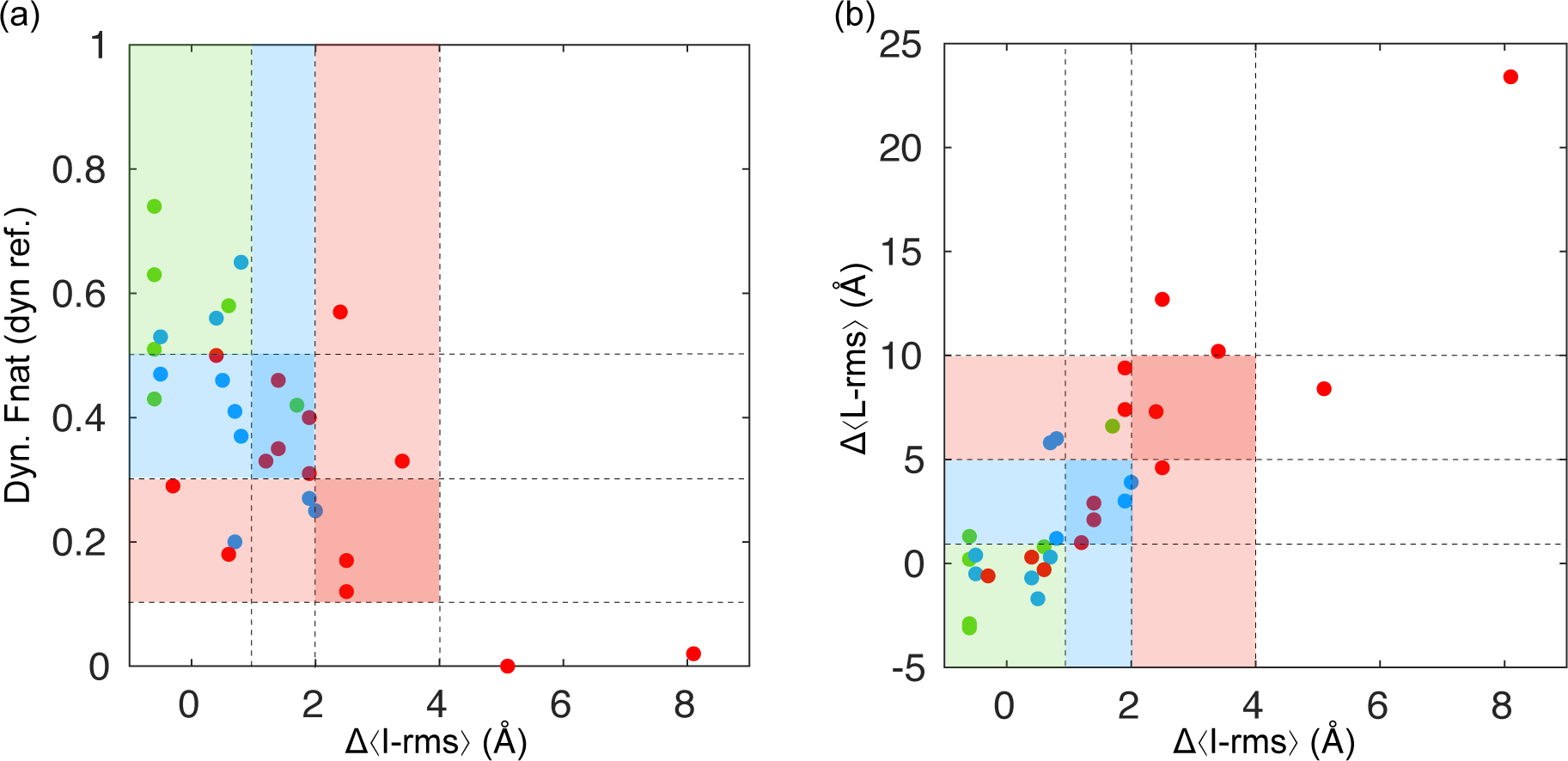
(a) Dynamic *f_nat_* as a function of Δ⟨I-rms⟩ (b) Δ⟨L-rms⟩ as a function of Δ⟨I- rms⟩. The areas corresponding to the new high, medium and acceptable categories (with dynamic criteria) are highlighted with green, blue and red backgrounds respectively.

**Table 2.** A summary of the studied models and their accuracy category, based on static or dynamic parameters.

| Model-Target | Static category | Dynamic category |
| --- | --- | --- |
| H604T41 | High | Medium |
| H1009T41 | High | High |
| M31T41 | Medium | High |
| M241T41 | Medium | Acceptable |
| M1149T41 | Medium | Acceptable |
| A50T41 | Acceptable | Medium |
| A61T41 | Acceptable | Acceptable |
| A302T41 | Acceptable | Medium |
| A380T41 | Acceptable | Medium |
| A490T41 | Acceptable | Acceptable |
| H190T29 | High | High |
| H1851T29 | High | Medium |
| M56T29 | Medium | High |
| M173T29 | Medium | Acceptable |
| M875T29 | Medium | Medium |
| M1818T29 | Acceptable | Medium |
| A608T29 | Acceptable | Acceptable |
| A1631T29 | Acceptable | Incorrect |
| A1909T29 | Acceptable | Acceptable |
| A2109T29 | Acceptable | Acceptable |
| H304T37 | High | High |
| H852T37 | High | High |
| M68T37 | Medium | Medium |
| M351T37 | Medium | Medium |
| M833T37 | Medium | High |
| A146T37 | Acceptable | High |
| A660T37 | Acceptable | Medium |
| A960T37 | Acceptable | Acceptable |
| A1022T37 | Acceptable | Incorrect |
| A1275T37 | Acceptable | Medium |

## 4. Conclusion

The assessment of the quality of protein complex structures predicted by docking approaches is traditionally done using criteria (*f_nat_*, L-rms and I-rms) that are based on a single reference experimental structure. However, this static view of the protein interface does not take into account its dynamic quality, which has been known for over a decade. In this study, we used all-atom classical Molecular Dynamics simulations to investigate the stability of the protein interface for three complexes that were previously used as targets in the CAPRI competition, and for each one of these targets, ten models retrieved from CAPRI Score-set 40 and distributed over the *High*, *Medium* and *Acceptable* accuracy categories. To assess the quality of these models from a dynamic perspective, we set up new dynamic definitions for the fraction of native contacts (*dyn- f_nat_*), the ligand- and the interface-rms (Δ⟨L-rms⟩ and Δ⟨I-rms⟩). These two parameters in particular take into account the stability of the reference protein interface along the MD trajectory. With this new set of criteria, more than 50% of the 30 models undergo a category change (either toward a better or a lower accuracy group) when reassessing their quality using dynamic information. Our results suggest that the static vision of the protein complex that goes with the reference crystallographic structure is probably not sufficient to assess the quality of a model produced by docking calculations, as it cannot guarantee that it will remain near this reference structure along time. One should note that obtaining an exhaustive sampling of the conformational space spanned by the protein-protein interface would require using several replica and performing longer trajectories. Such calculations would probably be too computationally costly to be systematically applied to the dozens of complex models that are usually retained after a docking procedure. However, our results show that MD simulations of reasonable length (and cost) are sufficient to retrieve relevant information regarding the stability of the model interfaces (and the stability of their accuracy), and compare it to the dynamics of the reference experimental interface. Altogether, MD simulations appear to be a necessary step following the docking and scoring procedures to ensure that the protein partners will actually remain assembled *happily ever after*.

## Acknowledgments

This work was supported by the “Initiative d’Excellence” program from the French State (Grant “DYNAMO”, ANR-11-LABX-0011-01). Simulations were performed using HPC resources from LBT/HPC

## Supporting Material

The values for the static and dynamic parameters defined in the *Material & Methods* section for the crystallographic structures and the 30 selected models are available as supporting information. SI also contains a list of the interface residues for the three targets using the static or the dynamic definitions, and additional data regarding the contacts formed at the protein interface during the MD simulations performed with the three targets crystallographic structures.

## References

1. Alberts B. The cell as a collection of protein machines: Preparing the next generation of molecular biologists. Cell. 1998;92(3):291–294.

2. Robinson CV, Sali A, Baumeister W. The molecular sociology of the cell. Nature. 2007;450(7172):973–982.

3. Ideker T, Sharan R. Protein networks in disease. Genome Res. 2008;18(4):644–652.

4. Wodak SJ, Pu S, Vlasblom J, Seraphin B. Challenges and rewards of interaction proteomics. Mol Cell Proteomics. 2009;8(1):3–18.

5. Wass MN, David A, Sternberg MJE. Challenges for the prediction of macromolecular interactions. Current Opinion in Structural Biology. 2011;21(3):382–390.

6. Garzon JI, Deng L, Murray D, Shapira S, Petrey D, Honig B. A computational interactome and functional annotation for the human proteome. eLife. 2016;5.

7. Keskin O, Tuncbag N, Gursoy A. Predicting Protein-Protein Interactions from the Molecular to the Proteome Level. Chem Rev. 2016;116(8):4884–4909.

8. Shoemaker BA, Panchenko AR. Deciphering protein-protein interactions. Part I. Experimental techniques and databases. PLoS Comput Biol. 2007;3(3):337–344.

9. Hein MY, Hubner NC, Poser I, et al. A human interactome in three quantitative dimensions organized by stoichiometries and abundances. Cell. 2015;163(3):712–723.

10. Berman HM, Battistuz T, Bhat TN, et al. The Protein Data Bank. Acta Crystallogr, sect D: Biol Crystallogr. 2002;58(Pt 6 No 1):899–907.

11. Bai XC, McMullan G, Scheres SH. How cryo-EM is revolutionizing structural biology. Trends Biochem Sci. 2015;40(1):49–57.

12. Cheng Y, Glaeser RM, Nogales E. How Cryo-EM Became so Hot. Cell. 2017;171(6):1229–1231.

13. Vamparys L, Laurent B, Carbone A, Sacquin-Mora S. Great interactions: How binding incorrect partners can teach us about protein recognition and function. Proteins. 2016;84(10):1408–1421.

14. Lagarde N, Carbone A, Sacquin-Mora S. Hidden partners: Using cross-docking calculations to predict binding sites for proteins with multiple interactions. Proteins. 2018;86(7):723–737.

15. Schweke H, Mucchielli MH, Sacquin-Mora S, Bei W, Lopes A. Protein Interaction Energy Landscapes are Shaped by Functional and also Non-functional Partners. J Mol Biol. 2020;432(4):1183–1198.

16. Shoemaker BA, Panchenko AR. Deciphering protein-protein interactions. Part II. Computational methods to predict protein and domain interaction partners. PLoS Comput Biol. 2007;3(4):595–601.

17. Kuzu G, Keskin O, Nussinov R, Gursoy A. Modeling protein assemblies in the proteome. Mol Cell Proteomics. 2014;13(3):887–896.

18. Vakser IA. Challenges in protein docking. Curr Opin Struct Biol. 2020;64:160–165.

19. Roel-Touris J, Jimenez-Garcia B, Bonvin A. Integrative modeling of membrane-associated protein assemblies. Nature communications. 2020;11(1):6210.

20. van Zundert GCP, Rodrigues J, Trellet M, et al. The HADDOCK2.2 Web Server: User-Friendly Integrative Modeling of Biomolecular Complexes. J Mol Biol. 2016;428(4):720–725.

21. Koukos PI, Bonvin A. Integrative Modelling of Biomolecular Complexes. J Mol Biol. 2020;432(9):2861–2881.

22. Rosell M, Fernandez-Recio J. Docking approaches for modeling multi-molecular assemblies. Curr Opin Struct Biol. 2020;64:59–65.

23. Mendez R, Leplae R, De Maria L, Wodak SJ. Assessment of blind predictions of protein-protein interactions: current status of docking methods. Proteins. 2003;52(1):51–67.

24. Lensink MF, Velankar S, Kryshtafovych A, et al. Prediction of homoprotein and heteroprotein complexes by protein docking and template-based modeling: A CASP-CAPRI experiment. Proteins. 2016;84 Suppl 1:323–348.

25. Lensink MF, Brysbaert G, Nadzirin N, et al. Blind prediction of homo- and hetero-protein complexes: The CASP13-CAPRI experiment. Proteins. 2019;87(12):1200–1221.

26. Mendez R, Leplae R, Lensink MF, Wodak SJ. Assessment of CAPRI predictions in rounds 3-5 shows progress in docking procedures. Proteins. 2005;60(2):150–169.

27. Lensink MF, Mendez R, Wodak SJ. Docking and scoring protein complexes: CAPRI 3rd edition. Proteins. 2007;69(4):704–718.

28. Fong JH, Shoemaker BA, Garbuzynskiy SO, Lobanov MY, Galzitskaya OV, Panchenko AR. Intrinsic disorder in protein interactions: insights from a comprehensive structural analysis. PLoS Comput Biol. 2009;5(3):e1000316.

29. Kuzu G, Gursoy A, Nussinov R, Keskin O. Exploiting conformational ensembles in modeling protein-protein interactions on the proteome scale. J Proteome Res. 2013;12(6):2641–2653.

30. Fuchs JE, Huber RG, Waldner BJ, et al. Dynamics Govern Specificity of a Protein-Protein Interface: Substrate Recognition by Thrombin. PLoS One. 2015;10(10):e0140713.

31. Visscher KM, Kastritis PL, Bonvin AM. Non-interacting surface solvation and dynamics in protein-protein interactions. Proteins. 2015;83(3):445–458.

32. Fuxreiter M. Fuzziness in Protein Interactions-A Historical Perspective. J Mol Biol. 2018;430(16):2278–2287.

33. Halakou F, Gursoy A, Keskin O. Embedding Alternative Conformations of Proteins in Protein-Protein Interaction Networks. Methods Mol Biol. 2020;2074:113–124.

34. Ozdemir ES, Nussinov R, Gursoy A, Keskin O. Developments in integrative modeling with dynamical interfaces. Curr Opin Struct Biol. 2019;56:11–17.

35. Kharche SA, Sengupta D. Dynamic protein interfaces and conformational landscapes of membrane protein complexes. Curr Opin Struct Biol. 2020;61:191–197.

36. Kurkcuoglu Z, Bonvin A. Pre- and post-docking sampling of conformational changes using ClustENM and HADDOCK for protein-protein and protein-DNA systems. Proteins. 2020;88(2):292–306.

37. van Wijk SJ, Melquiond AS, de Vries SJ, Timmers HT, Bonvin AM. Dynamic control of selectivity in the ubiquitination pathway revealed by an ASP to GLU substitution in an intra-molecular salt-bridge network. PLoS Comput Biol. 2012;8(11):e1002754.

38. Hou Q, Lensink MF, Heringa J, Feenstra KA. CLUB-MARTINI: Selecting Favourable Interactions amongst Available Candidates, a Coarse-Grained Simulation Approach to Scoring Docking Decoys. PLoS One. 2016;11(5):e0155251.

39. Nicoludis JM, Green AG, Walujkar S, et al. Interaction specificity of clustered protocadherins inferred from sequence covariation and structural analysis. Proc Natl Acad Sci U S A. 2019;116(36):17825–17830.

40. Lensink MF, Wodak SJ. Score_set: a CAPRI benchmark for scoring protein complexes. Proteins. 2014;82(11):3163–3169.

41. Meenan NA, Sharma A, Fleishman SJ, et al. The structural and energetic basis for high selectivity in a high-affinity protein-protein interaction. Proc Natl Acad Sci U S A. 2010;107(22):10080–10085.

42. Leulliot N, Chaillet M, Durand D, Ulryck N, Blondeau K, van Tilbeurgh H. Structure of the yeast tRNA m7G methylation complex. Structure. 2008;16(1):52–61.

43. Isabet T, Montagnac G, Regazzoni K, et al. The structural basis of Arf effector specificity: the crystal structure of ARF6 in a complex with JIP4. EMBO J. 2009;28(18):2835–2845.

44. Chakrabarti P, Janin J. Dissecting protein-protein recognition sites. Proteins-Structure Function and Genetics. 2002;47(3):334–343.

45. Kaminski GA, Friesner RA, Tirado-Rives J, Jorgensen WL. Evaluation and reparametrization of the OPLS-AA force field for proteins via comparison with accurate quantum chemical calculations on peptides. J Phys Chem B. 2001;105:6474–6487.

46. Jorgensen WL, Chandrasekhar J, Madura JD, Impey RW, Klein ML. Comparison of Simple Potential Functions for Simulating Liquid Water. J Chem Phys. 1983;79(2):926–935.

47. Bussi G, Donadio D, Parrinello M. Canonical sampling through velocity rescaling. J Chem Phys. 2007;126(1):014101.

48. Hess B, Bekker H, Berendsen HJC, Fraaije J. LINCS: A linear constraint solver for molecular simulations. J Comp Chem. 1997;18(12):1463–1472.

49. Essmann U, Perera L, Berkowitz ML, Darden T, Lee H, Pedersen LG. A SMOOTH PARTICLE MESH EWALD METHOD. J Chem Phys. 1995;103(19):8577–8593.

50. Parrinello M, Rahman A. Polymorphic transitions in single crystals: A new molecular dynamics method. J Appl Phys. 1981;52(12):7182–7190.

51. Boyer B, Danilowicz C, Prentiss M, Prevost C. Weaving DNA strands: structural insight on ATP hydrolysis in RecA-induced homologous recombination. Nucleic Acids Res. 2019;47(15):7798–7808.

52. Levy ED. A simple definition of structural regions in proteins and its use in analyzing interface evolution. J Mol Biol. 2010;403(4):660–670.

53. Humphrey W, Dalke A, Schulten K. VMD: visual molecular dynamics. J Mol Graph. 1996;14(1):33–38, 27-38.

