## Supplementary Information for "Moving pictures: Reassessing docking experiments with a dynamic view of protein interfaces"

**Table SI-1:** Summary of CAPRI assessment criteria for ranking predicted complexes (from Lensink et al., Proteins 2016)

| **Score** | $f_{nat}$ | | **L-rms** | | **I-rms** |
|---|---|---|---|---|---|
| **Incorrect** | < 0.1 | OR | ( > 10.0 | AND | > 4.0 ) |
| **Acceptable** | ≥ 0.3 | AND | > 5.0 | AND | > 2.0 |
| OR | $0.1 < f_{nat} < 0.3$ | AND | ( ≤ 10.0 | OR | ≤ 4.0 ) |
| **Medium** | ≥ 0.5 | AND | > 1.0 | AND | > 1.0 |
| OR | $0.3 \leq f_{nat} < 0.5$ | AND | ( ≤ 5.0 | OR | ≤ 2.0 ) |
| **High** | ≥ 0.5 | AND | ( ≤ 1.0 | OR | ≤ 1.0 ) |

**Table SI-2 :** values for the static and dynamic parameters defined in the *Material & Methods* section for the crystallographic structures and the 30 decoys.

| Decoy-Target | Initial Clashes | NstatCont | NdynCont | fnat/stat ref | dyn fnat/ stat ref | dyn fnat/ dyn ref | Lrmsd (in Å)/stat ref | Lrmsd (in Å) post equilibration | average Lrmsd (in Å) | min-max Lrmsd (in Å) | Irmsd (in Å)/stat ref | average Irmsd (in Å)/stat ref | min-max Irmsd (in Å)/ stat ref | average Irmsd (in Å)/ dyn ref | min-max Irmsd (in Å)/ dyn ref |
|---|---|---|---|---|---|---|---|---|---|---|---|---|---|---|---|
| 2wpt-T41 | 0 | 59 | 48 | 1.0 | 0.71 | 1.0 | 0.0 | 0.4 | 3.3 | 0.5-6.0 | 0.0 | 1.6 | 0.5-2.3 | 1.6 | 0.5-2.2 |
| H604-T41 | 13 | 60 | 47 | 0.88 | 0.41 | 0.42 | 1.4 | 0.5 | 9.9 | 1.5-15.3 | 0.6 | 3.3 | 0.8-4.1 | 3.3 | 0.8-4.2 |
| H1009-T41 | 18 | 55 | 42 | 0.78 | 0.49 | 0.58 | 2.6 | 0.4 | 4.1 | 1.9-6.4 | 0.8 | 2.2 | 0.9-2.9 | 2.2 | 0.9-2.9 |
| M31-T41 | 9 | 61 | 37 | 0.73 | 0.54 | 0.65 | 3.7 | 0.6 | 4.5 | 2.5-8.0 | 1.4 | 2.4 | 1.6-3.4 | 2.4 | 1.6-3.4 |
| M241-T41 | 5 | 67 | 56 | 0.70 | 0.24 | 0.27 | 6.6 | 0.6 | 6.3 | 4.2-9.7 | 1.7 | 3.5 | 2.9-3.9 | 3.5 | 2.9-4.0 |
| M1149-T41 | 7 | 64 | 60 | 0.68 | 0.27 | 0.25 | 6.6 | 0.6 | 7.2 | 3.0-12.0 | 1.6 | 3.6 | 3.0-4.1 | 3.6 | 2.9-4.1 |
| A50-T41 | 4 | 50 | 53 | 0.53 | 0.47 | 0.46 | 7.2 | 0.7 | 6.2 | 3.5-11.6 | 2.1 | 3.0 | 2.3-3.9 | 3.0 | 2.4-3.9 |
| A61-T41 | 3 | 45 | 23 | 0.46 | 0.22 | 0.17 | 6.6 | 0.7 | 7.9 | 3.3-11.0 | 2.1 | 4.1 | 2.3-5.3 | 4.0 | 2.3-5.2 |
| A302-T41 | 2 | 35 | 48 | 0.41 | 0.41 | 0.40 | 6.7 | 0.7 | 10.7 | 5.9-14.7 | 2.5 | 3.5 | 2.8-4.4 | 3.6 | 2.8-4.4 |
| A380-T41 | 3 | 46 | 49 | 0.53 | 0.31 | 0.31 | 7.1 | 0.6 | 12.7 | 6.9-16.9 | 2.2 | 3.5 | 2.2-4.2 | 3.5 | 2.2-4.2 |
| A490-T41 | 4 | 37 | 30 | 0.41 | 0.32 | 0.33 | 7.0 | 0.7 | 13.5 | 5.5-18.1 | 2.5 | 5.0 | 2.3-6.5 | 5.1 | 2.4-6.6 |
| 2vdu-T29 | 0 | 56 | 49 | 1.0 | 0.57 | 1.0 | 0.0 | 1.8 | 5.9 | 0.7-8.9 | 0.0 | 2.7 | 0.5-3.4 | 2.6 | 0.5-3.4 |
| H190T29 | 8 | 67 | 37 | 0.71 | 0.57 | 0.51 | 2.5 | 1.8 | 7.2 | 3.0-10.7 | 1.0 | 2.1 | 1.6-3.3 | 2.2 | 1.5-3.4 |
| H1851T29 | 10 | 71 | 29 | 0.76 | 0.50 | 0.43 | 3.2 | 1.8 | 6.1 | 2.9-10.0 | 1.0 | 2.1 | 1.5-3.1 | 2.1 | 1.5-3.2 |
| M56T29 | 4 | 63 | 48 | 0.69 | 0.68 | 0.53 | 3.9 | 1.8 | 5.4 | 2.4-10.3 | 1.1 | 2.2 | 1.7-3.3 | 2.1 | 1.5-3-4 |
| M173T29 | 3 | 46 | 26 | 0.45 | 0.23 | 0.20 | 4.1 | 1.8 | 11.7 | 4.6-18.4 | 1.8 | 3.4 | 2.3-4.9 | 3.4 | 2.3-4.8 |
| M875T29 | 3 | 40 | 37 | 0.37 | 0.45 | 0.37 | 6.2 | 1.8 | 11.9 | 5.7-17.0 | 1.5 | 3.5 | 2.1-4.4 | 3.5 | 2.0-4.4 |
| M1818T29 | 3 | 46 | 56 | 0.45 | 0.61 | 0.47 | 3.1 | 1.8 | 6.3 | 2.0-10.4 | 1.7 | 2.2 | 1.6-2.9 | 2.2 | 1.4-2.9 |
| A608T29 | 2 | 20 | 25 | 0.27 | 0.13 | 0.12 | 9.1 | 1.8 | 18.6 | 8.1-29.0 | 3.0 | 5.2 | 3.1-7.3 | 5.7 | 3.2-7.9 |
| A1631T29 | 3 | 60 | 40 | 0.14 | 0.0 | 0.0 | 9.5 | 0.5 | 14.3 | 8.7-20.2 | 4.4 | 7.8 | 4.7-8.6 | 8.0 | 4.8.07 |
| A1909T29 | 5 | 38 | 29 | 0.32 | 0.14 | 0.18 | 5.9 | 0.8 | 5.6 | 4.1-9.7 | 2.6 | 3.3 | 2.9-4.0 | 3.3 | 2.8-4.0 |
| A2109T29 | 5 | 40 | 29 | 0.40 | 0.34 | 0.29 | 5.8 | 1.7 | 5.3 | 2.8-11.5 | 2.4 | 2.4 | 1.8-3.4 | 2.4 | 1.8-3.4 |
| 2w83-T37 | 0 | 54 | 54 | 1.0 | 0.79 | 1.0 | 0.0 | 0.50 | 7.9 | 0.4-12.1 | 0.0 | 2.0 | 0.4-2.5 | 2.8 | 0.4-3.5 |
| H304T37 | 12 | 54 | 48 | 0.90 | 0.74 | 0.74 | 1.3 | 0.53 | 5 | 1.9-9.5 | 0.8 | 1.4 | 1.0-2.2 | 1.8 | 1.1-3.0 |
| H852T37 | 11 | 44 | 37 | 0.94 | 0.70 | 0.63 | 2.6 | 0.45 | 4.8 | 2.1-8.0 | 0.9 | 1.4 | 0.9-2.2 | 1.7 | 1.1-2.7 |
| M68T37 | 11 | 61 | 41 | 0.55 | 0.57 | 0.46 | 6.6 | 0.57 | 6.2 | 3.3-10.0 | 1.9 | 2.5 | 1.9-3.2 | 2.6 | 2.0-3.6 |
| M351T37 | 11 | 53 | 38 | 0.55 | 0.43 | 0.41 | 3.6 | 0.54 | 8.2 | 4.6-18.8 | 1.5 | 2.7 | 2.1-3.7 | 2.9 | 2.3-3.9 |
| M833T37 | 10 | 52 | 53 | 0.57 | 0.55 | 0.56 | 6.6 | 0.47 | 7.2 | 4.7-9.5 | 1.99 | 2.4 | 1.9-3.0 | 3.1 | 2.3-3.8 |
| A146T37 | 10 | 61 | 42 | 0.59 | 0.55 | 0.50 | 7.0 | 0.42 | 8.2 | 5.4-12.1 | 2.2 | 2.4 | 1.7-3.8 | 2.9 | 2.3-4.2 |
| A660T37 | 10 | 71 | 39 | 0.49 | 0.34 | 0.33 | 8.5 | 0.50 | 8.9 | 6.1-11.9 | 2.4 | 3.2 | 2.4-4.1 | 3.4 | 2.6-4.2 |
| A960T37 | 6 | 45 | 38 | 0.35 | 0.58 | 0.57 | 5.0 | 0.47 | 15.2 | 9.8-18.3 | 2.3 | 4.4 | 3.4-5.6 | 4.5 | 3.5-5.9 |
| A1022T37 | 10 | 51 | 3 | 0.14 | 0.02 | 0.02 | 9.7 | 0.53 | 31.3 | 19.5-45.0 | 4.4 | 10.1 | 6.4-14.1 | 10.7 | 6.9-14.9 |
| A1275T37 | 11 | 50 | 43 | 0.27 | 0.36 | 0.35 | 6.4 | 0.50 | 10.0 | 6.3-18.4 | 2.6 | 3.4 | 2.4-5.1 | 3.6 | 2.4-5.4 |

**Table SI-3:** List of interface residues for each target from the crystallographic structure (static interface) or the MD trajectory (dynamic interface)

| Target | Static interface | | Dynamic interface | |
|---|---|---|---|---|
| | Receptor residues | Ligand residues | Receptor residues | Ligand residues |
| **T41 (PDB 2WPT, chains B, A)** ***Colicin E9 DNase and Im2 immunity protein*** | 21-24, 26-27, 50-56, 58-59, 68-101, 123-124 | 18-42, 44-57, 60-67 | 21-26, 28, 50-52, 54-56, 58-59, 68-101, 123-124 | 19-58, 60-64, 67 |
| **T37 (PDB 2W83, chains A, CD)** ***Arf-effector complex*** | 12-17, 27, 30-31, 33-35, 44-54, 57-65, 68-82, 109-111 | 404-427, 430, 434, 1409-1411, 1413-1437, 1439 | 12-17, 30-31, 34-35, 39-54, 58-65, 68-81, 105, 108-111 | 398-399, 401-403, 405-431, 434, 1403, 1407, 1410-1411, 1413-1436, 1438-1439 |
| **T29 (PDB 2VDU, chains D,F)** ***Trm8/Trm82 tRNA guanine-N(7)-methyltransferase from S. cerevisiae*** | 168, 170-174, 188-198, 206-212, 215, 217-244, 253, 261, 265, 267, 269-274, 277 | 66-76, 127, 161-174, 195-209, 211, 227, 230-231, 234-238, 279-286 | 171-174, 189-197, 208-212, 215, 217-244, 259-263, 265, 267, 270-272, 281, 307 | 66-76, 127, 160-174, 197-209, 211, 227, 231, 234-238, 279-286 |

**Figure SI-1:** Cartoon representation of the target structures under study with the receptor chain in green, the ligand chain in its initial configuration (crystallographic structure) in purple/pink and the ligand chain by the end of the 100ns MD trajectory in blue/cyan.

(a) T41, (b) T29, (c) T37 (chain C in purple/blue and chain D in pink/cyan)

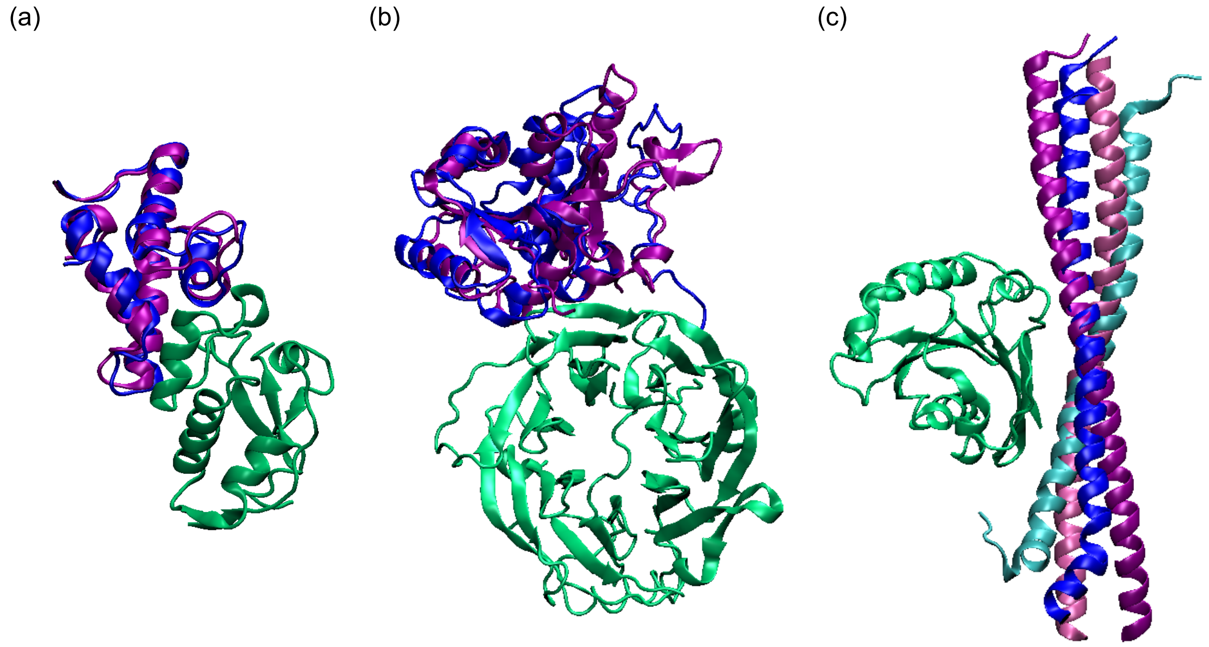

**Figure SI-2:** Target 41 (pdb 2wpt), time evolution during the MD simulation based on the crystallographic structure for:

(a) Number of contacts

(b) *fnat*, the fraction of native contacts from the reference structure that are still present in a given frame. The areas corresponding to the high, medium and acceptable categories are highlighted with green, orange and red backgrounds respectively.

(c) *fcont*, the fraction of contacts in a given frame that correspond to native contacts from the reference structure.

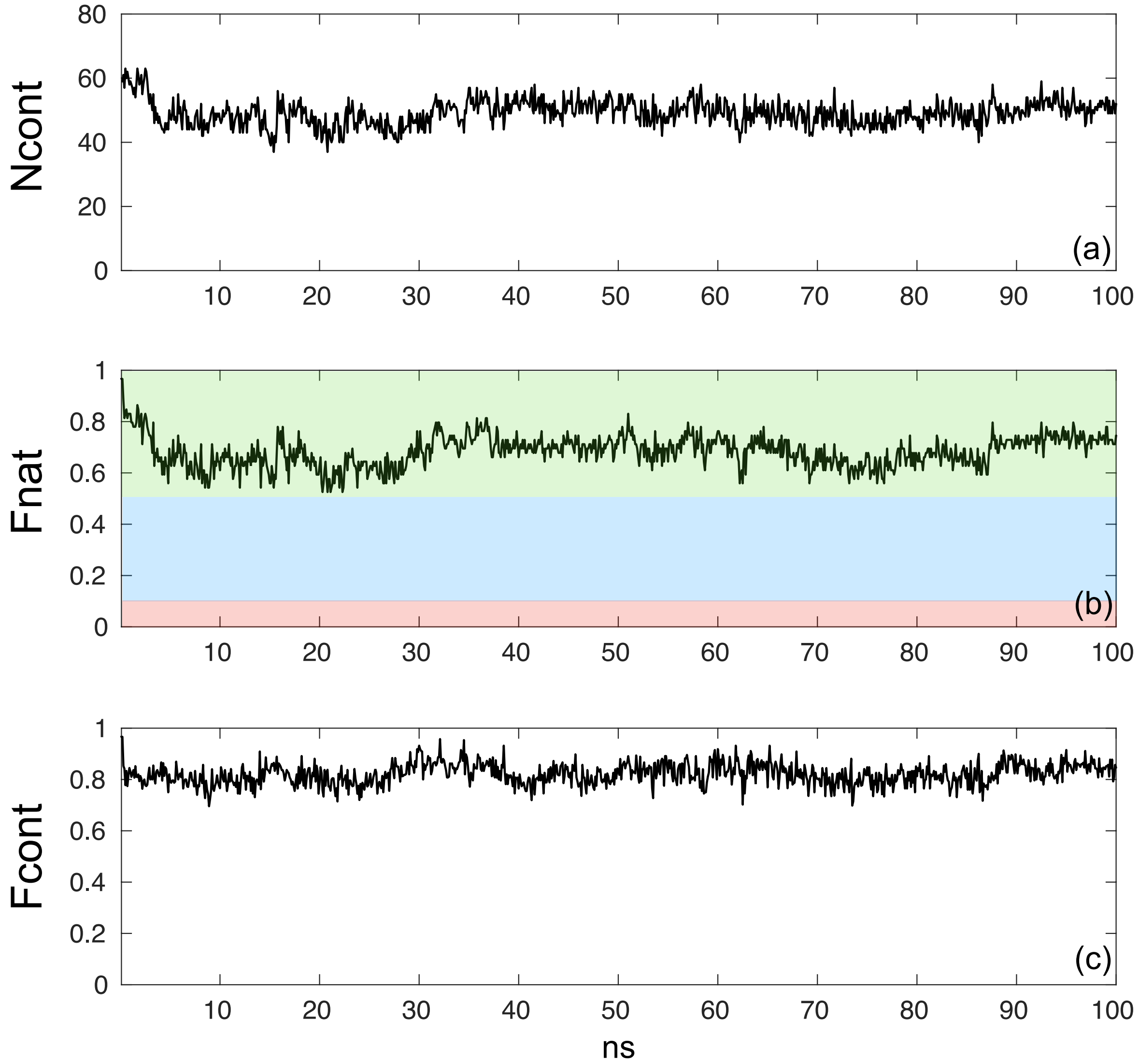

**Figure SI-3:** Target 29 (pdb 2vdu), time evolution during the MD simulation based on the crystallographic structure for:

(a) Number of contacts

(b) *fnat*, the fraction of native contacts from the reference structure that are still present in a given frame. The areas corresponding to the high, medium and acceptable categories are highlighted with green, orange and red backgrounds respectively.

(c) *fcont*, the fraction of contacts in a given frame that correspond to native contacts from the reference structure.

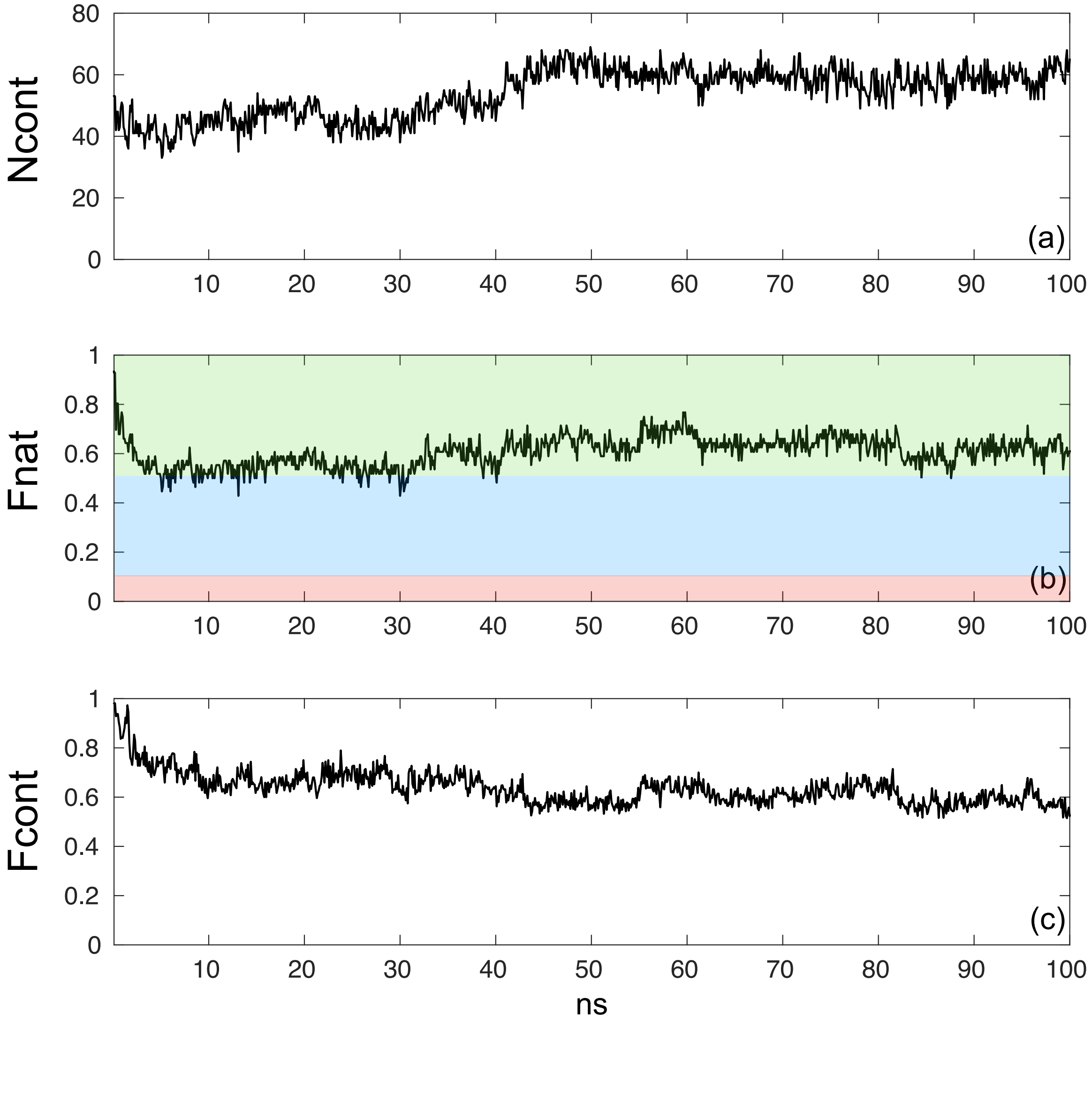

**Figure SI-4:** Target 37 (pdb 2w83), time evolution during the MD simulation based on the crystallographic structure for:

(a) Number of contacts

(b) *fnat*, the fraction of native contacts from the reference structure that are still present in a given frame. The areas corresponding to the high, medium and acceptable categories are highlighted with green, orange and red backgrounds respectively.

(c) *fcont*, the fraction of contacts in a given frame that correspond to native contacts from the reference structure.

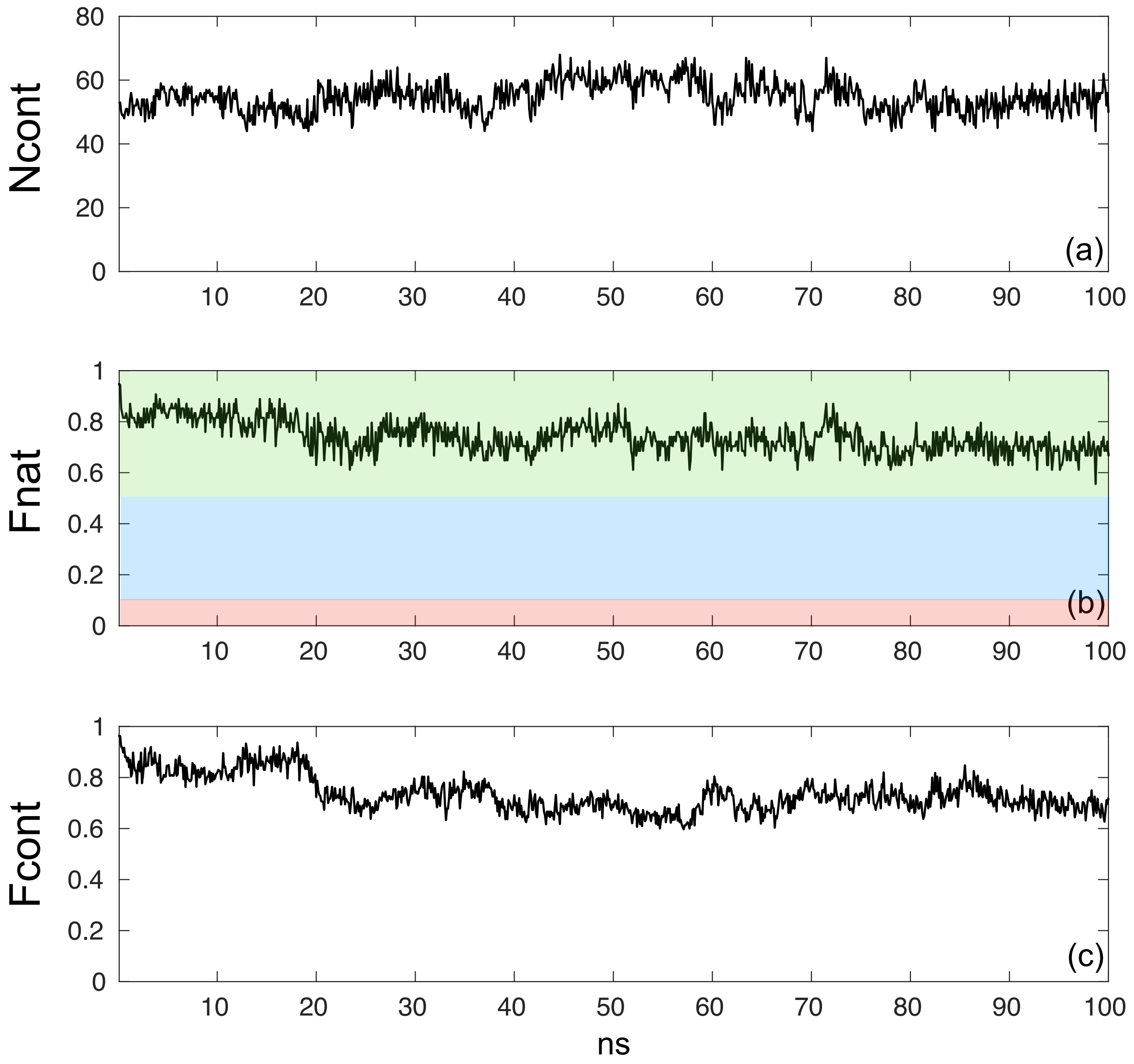

**Figure SI-5:** Number of residue pairs forming contacts during the trajectory (i.e. with heavy atoms less 5 Å from each other) as the function of their residence time for the three targets crystallographic structures. (a) T41-2wpt, (b) T29-2vdu, (c) T37-2w83

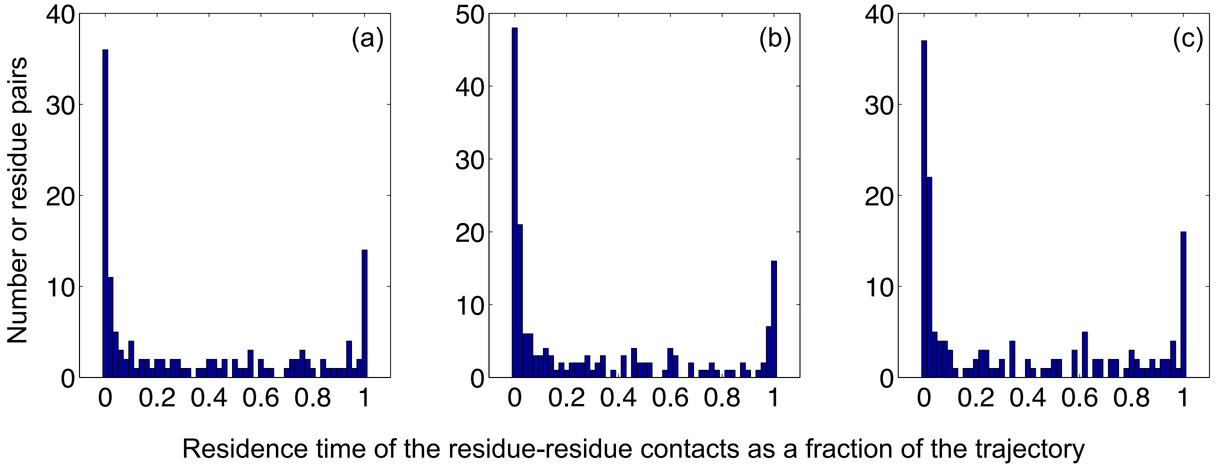

**Figure SI-6:** Post-equilibration L-rms as a function of the number clashes in the starting structure. Black dots: Crystallographic structures for the three targets. Green: High quality models. Orange: Medium quality models. Red: Acceptable quality models.

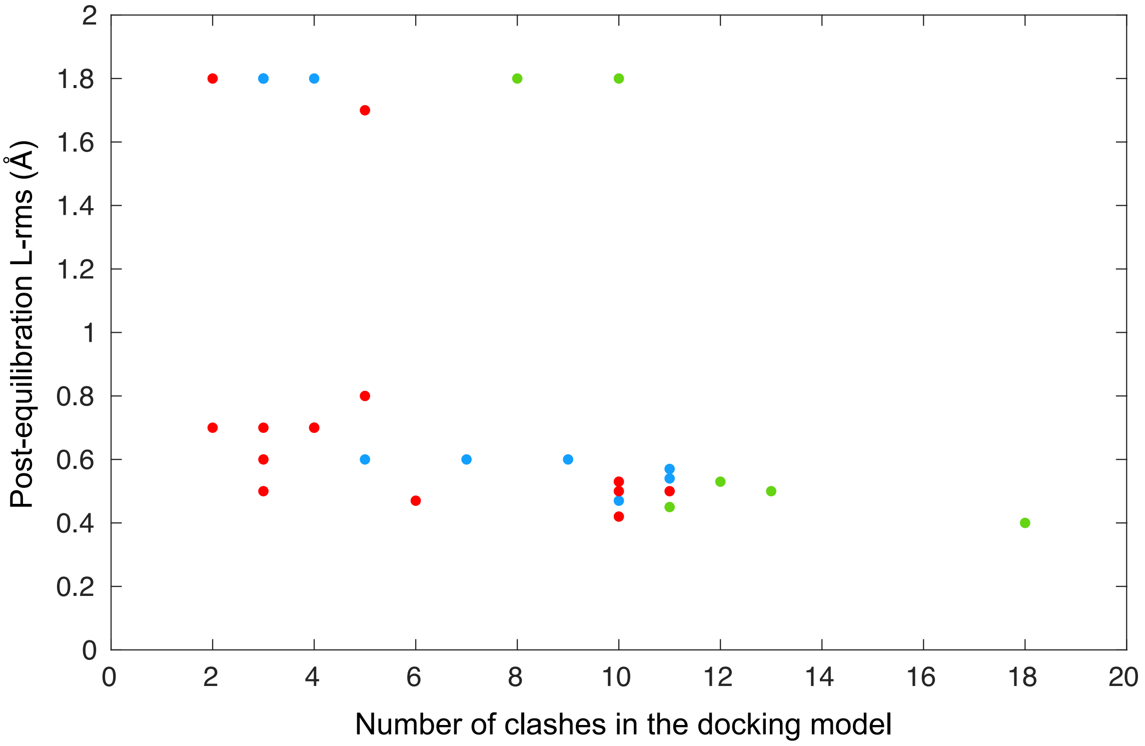
